# Synovial transcriptional clusters link cartilage degeneration to cell-type-specific gene expression in knee osteoarthritis

**DOI:** 10.64898/2026.04.21.719697

**Authors:** Michael R. Mazzucco, Bella Mehta, Jenelys Ruiz-Ortiz, Caryn Hale, Fuadur Omi, Purva Singh, Ruoxi Yuan, Samantha Lessard, Eun Kyung Song, Mengrui Zhang, Shady Younis, William H. Robinson, Daniel Ramirez, Edward DiCarlo, Wei Wang, Thomas Carroll, Jose Rodriguez, Peter Sculco, Xiaoshun Li, YiYuan Wu, Robert B. Darnell, Martin Lotz, Rachel E. Miller, Tristan Maerz, Anne-Marie Malfait, Accelerating Medicines Partnership: RA/SLE Network (AMP RA/SLE), The RE-JOIN Investigators, Miguel Otero, Dana E. Orange

## Abstract

**Objectives:** To identify synovial transcriptional clusters in human knee osteoarthritis (OA) and determine how these relate to synovial histologic features, cell-type-associated gene expression, and cartilage degeneration severity.

**Methods:** Bulk RNA sequencing (RNA-seq) of synovial tissue from n = 135 patients with knee OA was analyzed using consensus clustering. Clusters were compared by clinical and histologic features, including cartilage degeneration severity (OARSI score). Single-cell RNA-seq (n = 18) and spatial transcriptomics were used to relate cartilage degeneration-associated gene expression patterns to synovial cell populations.

**Results:** Four synovial transcriptional clusters that differed in synovial histologic features and cartilage degeneration severity were identified. Greater cartilage degeneration was associated with enrichment of lining fibroblast- and inflammatory myeloid-associated gene expression, whereas lesser cartilage degeneration was associated with enrichment of sublining fibroblast, endothelial, mural cell, and adipocyte-associated gene expression.

**Conclusions:** Human knee OA synovium segregates into transcriptional clusters associated with cartilage degeneration severity. Synovial transcriptional heterogeneity corresponds to cell-type-associated gene expression.

**Key messages:** *What is already known on this topic:*

- Osteoarthritis synovium exhibits marked histologic and molecular heterogeneity.
- Synovial inflammation detected by MRI correlates with cartilage degeneration and predicts progressive cartilage loss in knee OA.
- Prior transcriptomic studies have identified molecular subsets of OA synovium, but their relationship to cartilage degeneration severity remains unclear.

*What this study adds:*

- OA synovium segregates into four transcriptional clusters: Sublining (C1), Lymphomyeloid (C2), Myeloid (C3), and Major trauma (C4).
- Greater cartilage degeneration is associated with enrichment of inflammatory myeloid and lining fibroblast gene expression, whereas lesser degeneration is associated with enrichment of adipocyte, sublining fibroblast, endothelial, and mural cell–associated gene expression.

*How this study might affect research, practice or policy:*

- Provides a framework for a clinically relevant biological stratification of OA patients based on synovial molecular features.
- Informs future efforts to link synovial biology with OA prognosis, cartilage degeneration, treatment allocation, and development of targeted therapeutic strategies.

## Introduction

Knee osteoarthritis (OA), characterized by cartilage degeneration, pain, and functional impairment, affects more than 20% of adults aged 40 years and older worldwide[1]. The combination of high population prevalence, polygenic susceptibility, and late-life onset with cumulative environmental exposures predicts substantial heterogeneity in clinical and tissue-level phenotypes[2,3]. Resolving these clinically actionable biologically meaningful subsets remains a central challenge for the development of mechanism-based interventions in OA.

Among joint tissues, the synovium has emerged as a major source of heterogeneity in OA. The synovium is organized as a thin lining layer of PRG4+ fibroblasts and TREM2+ macrophages overlying a sublining fibroblast-vascular scaffold embedded in loose connective tissue and deeper adipose tissue[4–6]. Early histopathologic studies described distinct synovial patterns, including hyperplastic, inflammatory, fibrotic, and detritus-rich synoviopathy, distinguished by characteristic combinations of lining hyperplasia, immune infiltration, fibrosis, and cartilage or bone debris[4]. More recently, bulk transcriptomic profiling of human OA synovium has reinforced and extended these observations, demonstrating segregation of synovial tissue into two to four molecular subsets characterized by varying degrees of immune activation, cellular infiltration, and extracellular matrix remodeling[7–9]. Emerging work also suggests that OA is accompanied by remodeling of deeper adipose depots[10]. In end-stage disease, synovial adipose tissue exhibits inflammatory and fibrotic changes and reduced adipocyte-associated gene expression, particularly in patients with severe obesity[11].

Magnetic resonance imaging (MRI) studies in patients with knee OA have noted that synovial inflammation not only correlates with cartilage degeneration but also predicts subsequent cartilage loss[12–14], raising the possibility that synovial state may contribute to progressive cartilage degeneration, even when adjusting for baseline cartilage defects, meniscal tears, and extrusion. However, the molecular and cellular features of synovial tissue that relate to cartilage degeneration remain incompletely defined. Here, we analyzed synovial bulk and single-cell RNA sequencing (RNA-seq) with Xenium spatial transcriptomics, clinical data, and synovial and cartilage histopathology to define synovial transcriptional clusters and cell type-resolved gene expression associated with cartilage degeneration severity.

## Methods

### Patient Cohort

A total of 154 patients with knee osteoarthritis undergoing total knee arthroplasty (TKA) were enrolled with institutional review board approval and written informed consent. Synovial tissue from 135 patients was analyzed by bulk RNA sequencing, with additional samples used for single-cell RNA sequencing (n = 18) and spatial transcriptomics (n = 1). Patients met established clinical and radiographic criteria for knee osteoarthritis, and those with inflammatory or autoimmune rheumatic disease or other non-osteoarthritis indications for TKA were excluded. Additional cohort definitions and eligibility criteria are provided in the Supplementary Methods.

### Tissue Retrieval, Processing, and Histologic Scoring

Joint tissues were retrieved at the time of TKA and processed for pathologist-guided selection. Grossly diseased synovium was selected for analysis, with adjacent tissue preserved for histology and RNA isolation. Cartilage degeneration was assessed using the Osteoarthritis Research Society International (OARSI) grading system, and synovial histology was evaluated using established features of inflammation and tissue remodeling. Additional tissue processing and histologic scoring details are provided in the Supplementary Methods.

### Sequencing and Bioinformatic Analysis

Bulk synovial RNA was isolated and subjected to RNA sequencing, with sequencing quality control assessed using Picard. Transcript abundances were quantified with kallisto and summarized to gene-level estimates with tximport, followed by normalization for downstream analyses. Differential expression and pathway analyses were performed to define synovial transcriptional variation and its relationship to cartilage degeneration severity.

Single-cell RNA sequencing of cryopreserved synovial tissue was performed using the 10x Genomics platform, followed by alignment, quality control, doublet removal, ambient RNA correction, and comparison with external synovial reference datasets for clustering and annotation. Spatial transcriptomic profiling was performed on formalin-fixed paraffin-embedded synovium using the Xenium platform, with downstream analysis and cell-type annotation guided by single-cell–derived marker genes.

Detailed experimental procedures, preprocessing steps, and computational analyses are provided in the Supplementary Methods.

### Statistical Analysis

Statistical analyses were performed in R. Welch’s ANOVA or Kruskal–Wallis tests were used for continuous variables across multiple groups as appropriate, Wilcoxon tests were used for pairwise and one-sample comparisons, Fisher’s exact tests were used for categorical variables, and Spearman correlation coefficients were used for associations between continuous variables. P values were adjusted for multiple testing using the Benjamini–Hochberg method where applicable. Additional statistical details are provided in the Supplementary Methods.

### Patient and Public Involvement

Patients and/or the public were not involved in the design, conduct, reporting, or dissemination plans of this study.

## Results

### Consensus Clustering Identifies Four Transcriptionally Distinct Synovial Clusters in Knee Osteoarthritis

We performed unsupervised consensus clustering of bulk synovial RNA-seq from n = 135 patients undergoing TKA. Clustering used the 5,000 most variable genes (median absolute deviation), with 2,000 resampling iterations and 80% item subsampling. Across clustering solutions ranging from K = 2–10, four clusters emerged with clear block structure in the consensus matrix and diminishing gains in cumulative distribution function (CDF) area beyond K = 4, indicating stabilization of cluster structure at this resolution (ΔAUC at K = 4: 0.0346; ΔAUC at K = 5: 0.0216) (Figure 1A-C). At K = 4, samples segregated into four transcriptional clusters of sizes n = 39 (C1, 28.9%), n = 52 (C2, 37.8%), n = 36 (C3, 26.7%), and n = 9 (C4, 6.7%). PCA of the 5,000 most variable protein-coding genes demonstrated that consensus clusters aligned with the dominant axes of transcriptional variation, with separation visible along PC1 (19% variance explained) and PC2 (9% variance explained) (Figure 1D).

**Figure 1.**
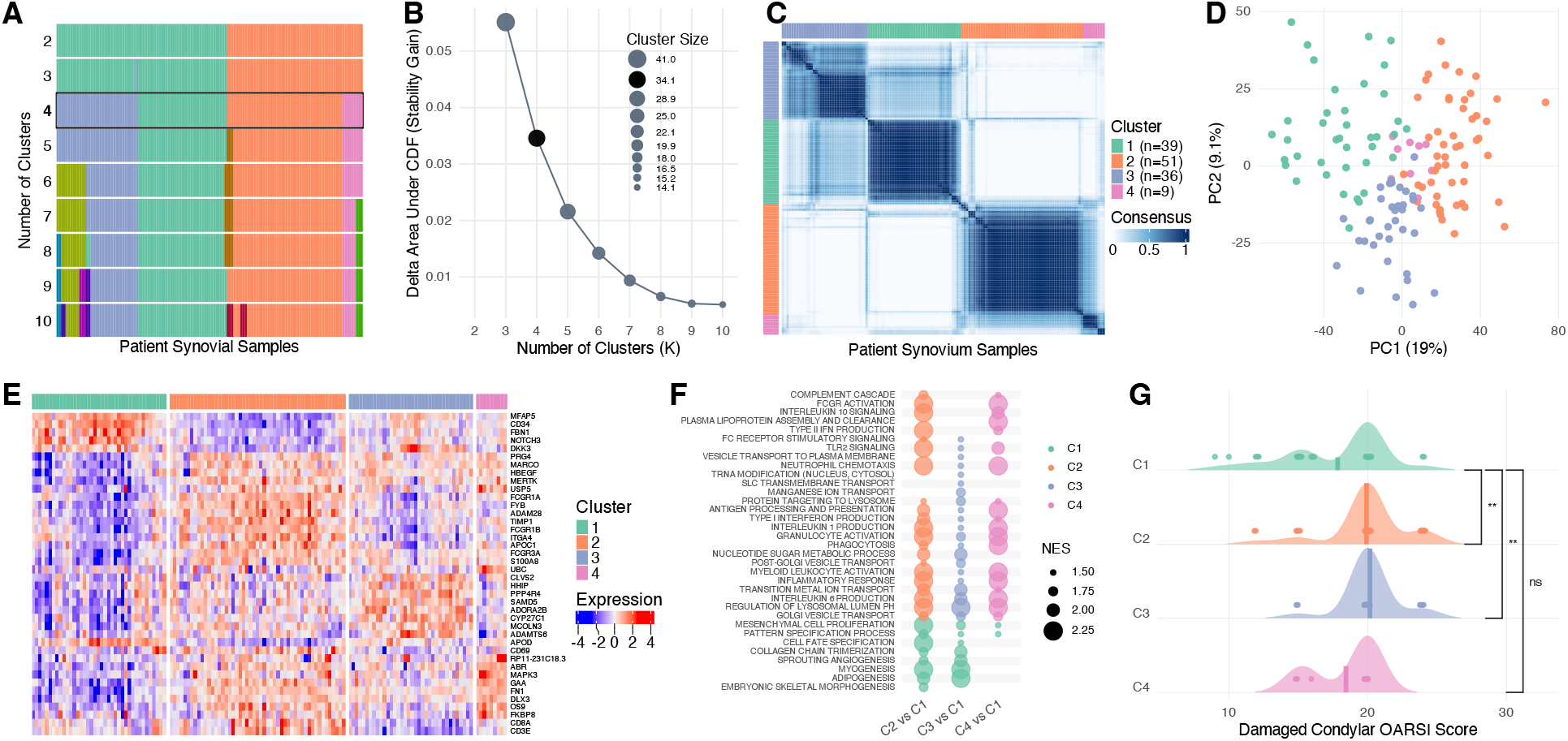
Consensus clustering identifies four robust synovial transcriptional clusters in knee osteoarthritis synovium. (A) Sample membership tracking across clustering solutions (K = 2–10). (B) Relative change in area under the consensus cumulative distribution function (ΔAUC) across K values. (C) Consensus matrix for K = 4 with cluster annotations; values indicate the fraction of resampling iterations in which each sample pair clustered together. (D) Principal component analysis (PCA) of the clustering gene set colored by cluster. (E) Heatmap of row-scaled expression for cluster-enriched marker genes, ordered by cluster. (F) Pathway enrichment across C2–C4 versus C1 contrasts (bubble size, normalized enrichment score; color, enrichment direction). (G) Damaged condylar OARSI scores by cluster with mean indicated; Wilcoxon tests versus C1. *P < 0.05, **P < 0.01, ***P < 0.001.

Pairwise differential expression analyses comparing each cluster to all remaining samples (cluster-versus-rest) using all expressed genes identified 4,033, 3,367, 2,059, and 1,281 differentially expressed genes (DEGs) in C1, C2, C3, and C4, respectively (FDR < 0.05). Genes selectively upregulated in each cluster were used to generate the cluster marker heatmap shown in Figure 1E, demonstrating substantial transcriptional heterogeneity across synovial samples. Complete differential expression results are provided in Supplemental Tables 6 and 7.

C1 expressed high levels of canonical synovial sublining fibroblast genes[6] including *FBN1, NOTCH3, MFAP5, APOD, CD34*, and *DKK3*, as well as YAP/TEAD and BMP/WNT family members. C2 showed enrichment of lymphocyte-associated genes such as *CD3E, CD8A, CD52*, and *CD69*. C3 exhibited increased expression of lining fibroblast and macrophage-associated genes including *PRG4, HBEGF, MARCO, S100A8, MERTK*, and *FCGR3A*. C4, the smallest cluster, was characterized by expression of genes involved in unfolded protein stress response and autophagy, including *OS9, FKBP8, UBC, USP5*, and *USP11*.

We next performed pathway enrichment analysis comparing each cluster to C1, which represented a stromal-dominant transcriptional state. C1 was enriched in mesenchymal proliferation, collagen organization, angiogenesis, adipogenesis, and myogenesis pathways, whereas C2 showed the strongest enrichment of immune-associated pathways, including antigen processing and presentation, B-cell receptor signaling, Fc receptor signaling, and interferon responses. Compared with C1, C3 and C4 were also enriched for inflammatory pathways, though to a lesser degree than C2 (Figure 1F). Complete pathway enrichment results are provided in Supplemental Tables 8 and 9. Together, these analyses indicate that bulk synovial transcriptomes in knee OA segregate into four transcriptionally distinct clusters characterized by varying degrees of inflammatory and fibroblast-associated gene expression.

Histological cartilage degeneration severity differed modestly but significantly across transcriptional clusters (Figure 1G). Damaged condylar OARSI scores were tightly distributed at the upper end of the scale across all groups, with median scores of 20 in C1 (n = 39), C2 (n = 50), C3 (n = 35), and C4 (n = 9). Relative to C1, OARSI scores were higher in C2 (Wilcoxon P = 0.0029) and C3 (P = 0.0015), whereas C4 did not differ significantly from C1 (P = 0.756).

### Clinical, Radiographic and Histologic Features Vary Across Synovial Transcriptional Clusters

We next examined whether synovial transcriptional clusters differed with respect to patient characteristics, disease severity, and synovial tissue-level pathology (Tables 1 and 2). Demographic variables including age, sex, body mass index, and duration of symptoms or diagnosis were similar across clusters, as were patient-reported pain scores and systemic inflammatory markers (ESR and CRP).

**Table 1.** Clinical characteristics across transcriptional clusters in knee osteoarthritis. Clinical and demographic characteristics of individuals with knee osteoarthritis stratified by synovial transcriptional cluster are shown. Continuous variables are reported as mean (SD) when approximately normally distributed across clusters, as assessed by the Shapiro-Wilk test, and as median (p25, p75) otherwise; categorical variables are reported as n (%). P values were calculated using Welch’s one-way ANOVA for normally distributed continuous variables, Kruskal-Wallis tests otherwise, and Fisher’s exact tests for categorical variables. Statistically significant values (P < 0.05) are shown in bold.

| Variable | Sublining (C1)<br>N = 39<sup>1</sup> | Lymphomyeloid (C2)<br>N = 51<sup>1</sup> | Myeloid (C3)<br>N = 36<sup>1</sup> | Major trauma (C4)<br>N = 9<sup>1</sup> | p-value<sup>2</sup> |
| --- | --- | --- | --- | --- | --- |
| Age at Baseline | 66.2 (60.6, 70.3) | 66.7 (61.5, 70.8) | 63.6 (59.4, 68.8) | 69.7 (62.6, 70.2) | 0.5 |
| Gender |  |  |  |  | 0.3 |
| Female | 24 (62%) | 34 (67%) | 21 (58%) | 3 (33%) |  |
| Male | 15 (38%) | 17 (33%) | 15 (42%) | 6 (67%) |  |
| Body Mass Index (BMI) | 29.6 (28.2, 31.3) | 29.5 (26.6, 37.3) | 30.3 (26.9, 33.4) | 27.3 (27.0, 28.6) | 0.5 |
| Race |  |  |  |  | 0.2 |
| White | 34 (87%) | 36 (71%) | 30 (83%) | 9 (100%) |  |
| Black or African American | 0 (0%) | 6 (12%) | 1 (3%) | 0 (0%) |  |
| Asian | 2 (5%) | 8 (16%) | 3 (8%) | 0 (0%) |  |
| American Indian or Alaska Native | 1 (3%) | 0 (0%) | 0 (0%) | 0 (0%) |  |
| Native Hawaiian or Other Pacific Islander | 1 (3%) | 0 (0%) | 0 (0%) | 0 (0%) |  |
| More than one race | 0 (0%) | 1 (2%) | 1 (3%) | 0 (0%) |  |
| Other | 1 (3%) | 0 (0%) | 1 (3%) | 0 (0%) |  |
| Diagnosis Duration (Years) | 9.4 (3.6, 15.0) | 5.8 (3.2, 14.4) | 6.7 (3.5, 16.2) | 7.4 (3.3, 11.1) | 0.8 |
| Symptom Duration (Years) | 10.7 (5.3, 15.7) | 10.5 (5.7, 19.6) | 13.9 (5.3, 30.3) | 9.6 (7.2, 10.7) | 0.8 |
| Duration of Pain (Years) | 9.5 (3.6, 12.7) | 6.3 (4.5, 15.1) | 12.4 (5.0, 23.3) | 10.7 (7.5, 20.5) | 0.11 |
| Kellgren–Lawrence Score | 3.0 (3.0, 4.0) | 4.0 (3.0, 4.0) | 3.0 (3.0, 4.0) | 3.0 (3.0, 4.0) | **0.045** |
| Control Condylar OARSI Score | 5.0 (3.0, 9.0) | 8.0 (3.0, 12.0) | 4.0 (3.0, 12.0) | 6.0 (4.0, 9.0) | 0.5 |
| Damaged Condylar OARSI Score | 20.0 (15.0, 20.0) | 20.0 (20.0, 20.0) | 20.0 (20.0, 20.0) | 20.0 (16.0, 20.0) | **0.002** |
| KOOS Pain Score | 51.1 (16.1) | 53.4 (15.2) | 53.5 (21.6) | 59.6 (13.7) | 0.5 |
| ESR | 11.0 (6.5, 23.0) | 16.0 (8.0, 22.0) | 16.5 (5.0, 29.5) | 8.0 (5.0, 11.0) | 0.2 |
| CRP | 1.6 (0.6, 3.4) | 1.7 (0.8, 4.5) | 1.3 (0.9, 2.9) | 0.8 (0.7, 1.1) | 0.11 |
| Any Prior Knee Injury | 27 (69%) | 28 (55%) | 22 (61%) | 8 (89%) | 0.2 |
| ACL Tear | 2 (5%) | 2 (4%) | 5 (14%) | 2 (25%) | 0.10 |
| Meniscal Tear | 21 (54%) | 22 (43%) | 16 (44%) | 8 (89%) | 0.067 |
| Major Trauma (Operated Joint) | 4 (10%) | 2 (4%) | 5 (14%) | 3 (38%) | **0.037** |
| Fracture (Leg) | 3 (8%) | 3 (6%) | 1 (3%) | 1 (13%) | 0.5 |
| Bleeding Into Joint | 0 (0%) | 0 (0%) | 2 (6%) | 0 (0%) | 0.2 |
| Other Fractures | 8 (21%) | 12 (24%) | 9 (25%) | 2 (25%) | >0.9 |
| Injury Score (0–6) | 1.0 (0.0, 2.0) | 1.0 (0.0, 1.0) | 1.0 (0.0, 2.0) | 2.0 (1.0, 3.0) | 0.075 |
| <sup>1</sup> Median (Q1, Q3); n (%); Mean (SD) |  |  |  |  |  |
| <sup>2</sup> Kruskal-Wallis rank sum test; One-way analysis of means (not assuming equal variances) |  |  |  |  |  |

**Table 2.** Histopathologic synovial features across transcriptional clusters. Synovial histopathologic characteristics of individuals with knee osteoarthritis stratified by synovial transcriptional cluster are shown. Continuous variables are reported as mean (SD) when approximately normally distributed across clusters, as assessed by the Shapiro-Wilk test, and as median (p25, p75) otherwise; categorical variables are reported as n (%). P values were calculated using Welch’s one-way ANOVA or Kruskal-Wallis tests for continuous variables, as appropriate, and Fisher’s exact tests for categorical variables. Statistically significant values (P < 0.05) are shown in bold.

| Variable | Sublining (C1)<br>N = 39<sup>1</sup> | Lymphomyeloid (C2)<br>N = 51<sup>1</sup> | Myeloid (C3)<br>N = 36<sup>1</sup> | Major trauma (C4)<br>N = 9<sup>1</sup> | p-value<sup>2</sup> |
| --- | --- | --- | --- | --- | --- |
| Inflammation (cell density $\times 10^3$) | 2.7 (2.1, 3.0) | 2.8 (2.4, 3.2) | 2.5 (1.9, 2.7) | 2.4 (2.2, 3.0) | **0.005** |
| Synovial lining inflammation |  |  |  |  | **0.001** |
| Low | 39 (100%) | 42 (82%) | 36 (100%) | 9 (100%) |  |
| High | 0 (0%) | 9 (18%) | 0 (0%) | 0 (0%) |  |
| Vascularity |  |  |  |  | **0.036** |
| Normal | 29 (74%) | 28 (55%) | 17 (47%) | 8 (89%) |  |
| Slight | 10 (26%) | 22 (43%) | 19 (53%) | 1 (11%) |  |
| Marked | 0 (0%) | 1 (2%) | 0 (0%) | 0 (0%) |  |
| Fibrosis |  |  |  |  | 0.2 |
| None | 2 (5%) | 2 (4%) | 2 (6%) | 0 (0%) |  |
| Focal | 25 (64%) | 33 (65%) | 14 (39%) | 6 (67%) |  |
| Widespread/Band-like | 12 (31%) | 16 (31%) | 20 (56%) | 3 (33%) |  |
| Synovial mucoid change |  |  |  |  | 0.12 |
| None | 2 (5%) | 0 (0%) | 1 (3%) | 0 (0%) |  |
| Slight | 17 (44%) | 21 (41%) | 20 (56%) | 1 (11%) |  |
| Moderate | 15 (38%) | 18 (35%) | 9 (25%) | 7 (78%) |  |
| Marked/Myxomatous | 5 (13%) | 12 (24%) | 6 (17%) | 1 (11%) |  |
| Synovial lining hyperplasia |  |  |  |  | 0.086 |
| Normal | 2 (5%) | 3 (6%) | 3 (8%) | 1 (11%) |  |
| 2–3 cells | 31 (79%) | 27 (53%) | 26 (72%) | 4 (44%) |  |
| 3–4 cells | 6 (15%) | 20 (39%) | 6 (17%) | 4 (44%) |  |
| >4 cells | 0 (0%) | 1 (2%) | 1 (3%) | 0 (0%) |  |
| Detritus |  |  |  |  | 0.15 |
| Absent | 27 (69%) | 24 (47%) | 22 (61%) | 4 (44%) |  |
| Small | 12 (31%) | 27 (53%) | 14 (39%) | 5 (56%) |  |
| Large | 0 (0%) | 0 (0%) | 0 (0%) | 0 (0%) |  |
| Plasma cell inflammation |  |  |  |  | **0.008** |
| <10% plasma | 33 (85%) | 37 (73%) | 35 (97%) | 9 (100%) |  |
| <50% plasma | 6 (15%) | 14 (27%) | 1 (3%) | 0 (0%) |  |
| >50% plasma | 0 (0%) | 0 (0%) | 0 (0%) | 0 (0%) |  |
| Binucleate plasma cells |  |  |  |  | **0.015** |
| None | 35 (90%) | 41 (80%) | 36 (100%) | 9 (100%) |  |
| Present | 4 (10%) | 10 (20%) | 0 (0%) | 0 (0%) |  |
| Russell bodies |  |  |  |  | 0.4 |
| None | 36 (92%) | 43 (84%) | 34 (94%) | 9 (100%) |  |
| Present | 3 (8%) | 8 (16%) | 2 (6%) | 0 (0%) |  |
| <sup>1</sup> Median (Q1, Q3); n (%) |  |  |  |  |  |
| <sup>2</sup> Kruskal-Wallis rank sum test |  |  |  |  |  |

In contrast, clusters differed in measures of radiographic and cartilage degeneration severity. Kellgren–Lawrence grades varied modestly across clusters (P = 0.045), with higher grades observed more frequently in C2 (median 4 [3–4]) compared with C1, C3, and C4 (each median 3 [3–4]). Pathologist-assessed cartilage degeneration also differed significantly across clusters (P = 0.002). Although damaged-condyle OARSI medians were most commonly found at 20 across clusters, reflecting advanced disease in this surgical cohort, the distribution was shifted lower in C1 (20 [15–20]) compared with C2 and C3 (both 20 [20–20]) and C4 (20 [16–20]), indicating relatively less severe cartilage degeneration in C1 overall. In contrast, control condyle OARSI scores did not differ across clusters (P = 0.5), suggesting cluster differences primarily reflect variation in degeneration at the affected site.

C4 was distinguished by clinical history, with a higher proportion of patients reporting prior major trauma involving the operative knee (38% vs 4–14% in other clusters; P = 0.037) and a similar trend toward increased prior meniscal tears (89% vs 43–54%; P = 0.067), accompanied by higher injury scores overall (2 [1–3] vs 1 [0–2]; P = 0.075).

Similarly, histologic measures of synovial inflammation also varied across clusters. Computer vision-derived synovial cell density differed across clusters and was modestly higher in C2 (2.8 ×10^3^ cells [2.4–3.2]) compared with C1 (2.7 [2.1–3.0]), C3 (2.5 [1.9–2.7]), and C4 (2.4 [2.2–3.0]; P = 0.005). Synovial lining inflammation and plasma cell-associated features were also enriched in C2, including higher lining inflammation scores (P = 0.001), greater plasma cell inflammation (P = 0.008), and increased presence of binucleate plasma cells (P = 0.015) (Table 2).Vascularity also differed across clusters (P = 0.037), with a greater proportion of samples showing slight or marked vascularity in C2 and C3 relative to C1 and C4, consistent with increased vascular remodeling in clusters associated with more severe cartilage degeneration. Other histologic features, including fibrosis, mucoid change, synovial lining hyperplasia, detritus, and Russell bodies, did not significantly differ across clusters.

To place subsequent single-cell analyses in context, we next compared clinical characteristics between patients included in the bulk RNA sequencing and single-cell cohorts and observed broadly similar demographic and disease features between cohorts (Supplemental Table 5).

### Single-cell RNA Sequencing Identifies Cell-type Gene Expression Patterns Underlying Synovial Transcriptional Clusters

To further interpret the transcriptional variation observed in OA synovial bulk RNA-seq, we used scRNA-seq data generated from synovium obtained from n = 18 additional patients. Single-cell analysis identified eight major cell populations, including T cells, B cells, plasma cells, myeloid cells, endothelial cells, lining fibroblasts, sublining fibroblasts, and ACTA2-positive mural cells (Figure 2A). We then defined high-confidence cell type-specific marker gene sets using a Wilcoxon rank-sum framework with stringent enrichment and prevalence thresholds (log2FC ≥ 0.5, expressed in ≥ 40% of target cells, and detected in a ≥ 10% higher fraction of cells than all other cells) (Figure 2B; Supplemental Table 4).

**Figure 2.**
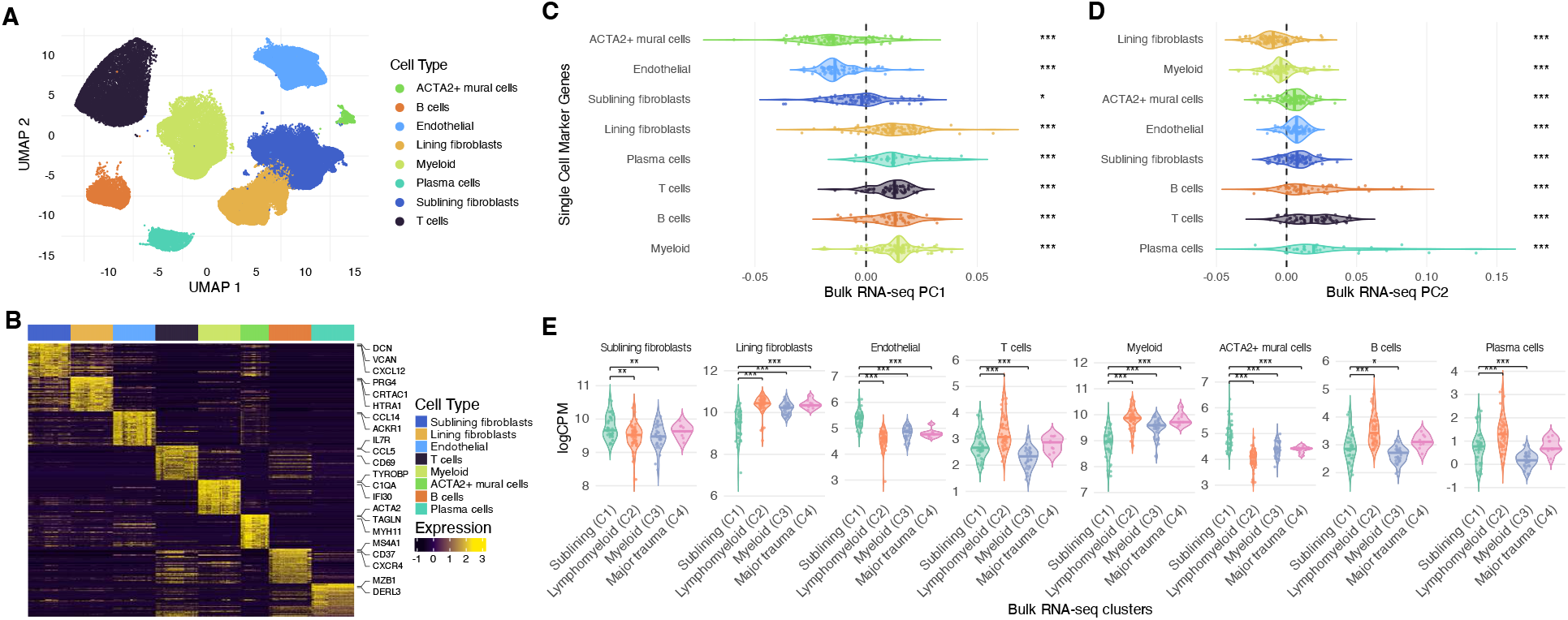
Single-cell RNA sequencing identifies cell-type gene expression patterns underlying synovial transcriptional clusters. (A) UMAP of synovial single cells from osteoarthritis samples colored by cell type. (B) Heatmap of cell-type–specific marker genes derived from scRNA-seq. (C,D) Distributions of single-cell marker gene loadings on bulk RNA-seq principal components PC1 and PC2. Significance was assessed using a one-sample Wilcoxon test against zero. (E) Mean bulk expression of scRNA-seq-derived cell-type marker genes (logCPM; top markers per cell type) across transcriptional clusters. Significance was tested using one-way ANOVA with Dunnett’s post hoc test comparing each cluster to Sublining (C1). *P < 0.05, **P < 0.01, ***P < 0.001.

We next assessed whether scRNA-seq-derived cell-type marker genes were enriched among genes contributing to PC1 and PC2 in the bulk RNA-seq PCA (Figures 2C-D). Myeloid, lining fibroblast, and lymphoid marker genes contributed strongly to positive PC1 loadings, whereas ACTA2-positive mural cell, endothelial cell, and sublining fibroblast markers were associated with lower PC1 loadings. In contrast, PC2 loadings were highest for infiltrating lymphocyte populations, including T cells, B cells, and plasma cells, and lowest for lining fibroblast and myeloid gene expression. Together, these findings indicate that the dominant axis of synovial transcriptional variation (PC1) reflects a gradient from stromal-dominant to inflammatory cell-associated gene expression, while PC2 primarily captures variation in lymphocyte-associated gene expression across samples.

Mean bulk expression of scRNA-seq-derived cell-type marker genes differed significantly across transcriptional clusters (one-way ANOVA with Dunnett’s post-hoc test versus C1) (Figure 2E). Full marker gene results for each synovial cell type are provided in Supplemental Table 4. On the basis of these cell-type markers, clusters are hereafter referred to as Sublining (C1), Lymphomyeloid (C2), Myeloid (C3), and Major trauma (C4). The Sublining (C1) cluster, which was relatively low along PC1 and enriched for mesenchymal proliferation and angiogenesis pathways (Figure 1F), showed higher expression of sublining fibroblast, endothelial, and mural cell marker genes compared with the other clusters. Relative to Sublining (C1), the Lymphomyeloid cluster (C2) demonstrated increased expression of T-cell, B-cell, plasma cell, myeloid, and lining fibroblast marker genes. The Myeloid cluster (C3) exhibited increased lining fibroblast and myeloid marker gene expression, but relatively lower T-cell and plasma cell marker gene expression compared with C2.

Both Lymphomyeloid (C2) and Myeloid (C3) clusters were associated with increased cartilage degeneration scores (Table 2), despite marked differences in adaptive immune cell-associated genes, suggesting that synovial lining fibroblast and myeloid gene expression, rather than lymphocyte-associated gene expression alone, are more closely linked to cartilage degeneration in OA.

### Synovial Gene Expression Associated with Cartilage Degeneration

The observation that OARSI cartilage scores varied across clusters suggested synovial gene expression associated with cartilage degeneration. We compared bulk synovial RNA-seq profiles between samples with high cartilage degeneration (D-OARSI score ≥ 20) and those with lower degeneration, irrespective of transcriptional cluster. Full statistical outputs supporting these analyses are provided in Supplemental Table 2. Differential expression analysis identified n = 337 genes differentially expressed between high and low cartilage degeneration groups, including n = 135 genes with increased expression and n = 202 genes with decreased expression in high-degeneration samples (Figure 3A). The complete differential expression results are provided in Supplemental Table 2.

**Figure 3.**
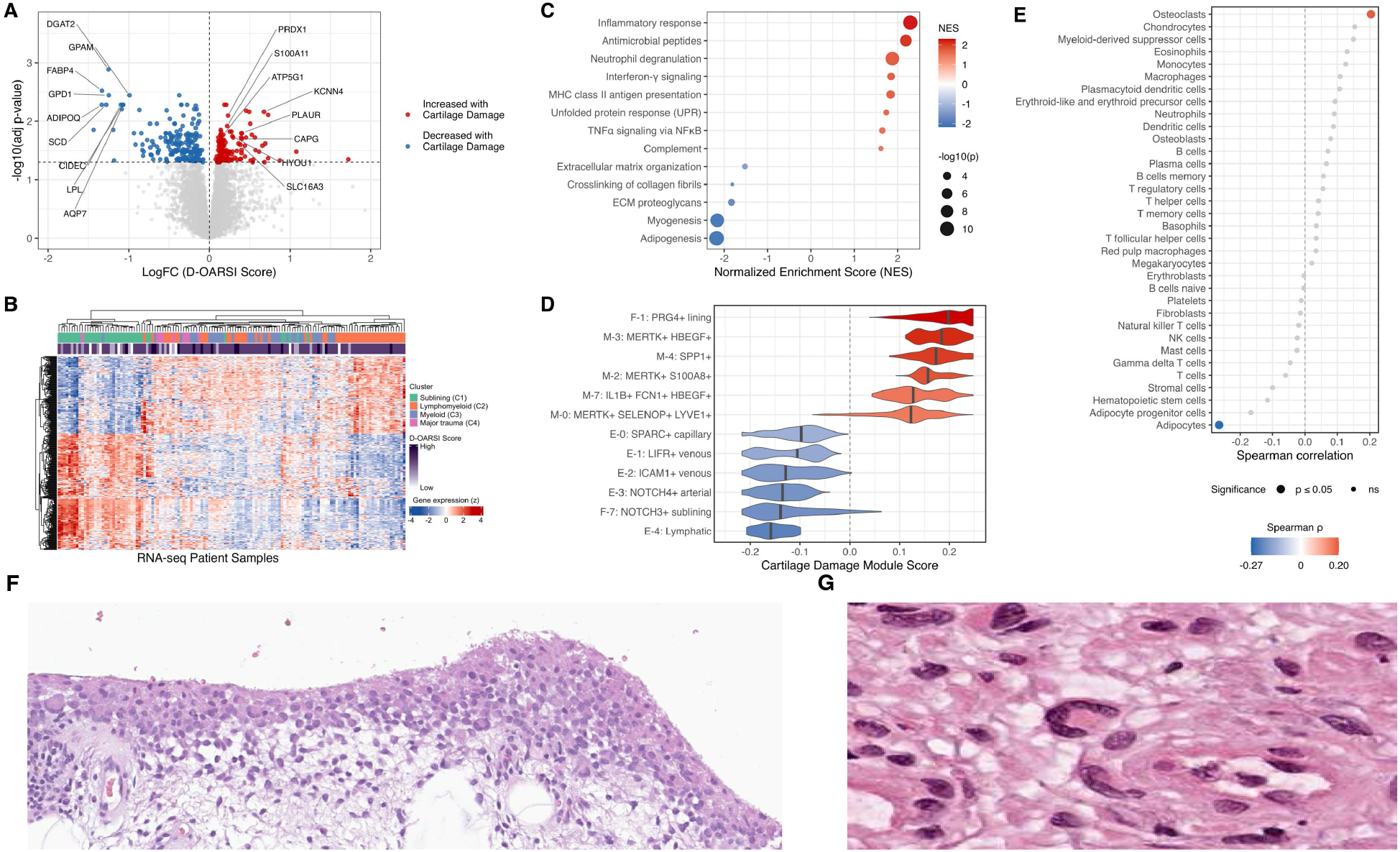
Synovial gene expression associated with cartilage degeneration in osteoarthritis. (A) Volcano plot of differential gene expression in OA knee synovium bulk RNA-seq comparing high cartilage degeneration (D-OARSI ≥ 20) versus low degeneration, with selected genes annotated. (B) Heatmap representation of significantly associated genes (row Z-scored) across synovial RNA-seq patient samples, annotated by bulk transcriptional cluster and D-OARSI score. (C) Pathway enrichment by FGSEA using ranked differential expression statistics (Hallmark and Reactome; bubble size, −log10(P); color, normalized enrichment score (NES)). (D) Synovial single-cell mapping of a cartilage degeneration–associated module score using KNN-smoothed per-cell scores across selected broad cell types and subtypes (violin plots; median indicated). (E) Spearman correlations between Panglao-derived gene set GSVA scores and D-OARSI score (dot position, Spearman ρ; dot size, −log10(P); color indicates significant correlations; non-significant sets shown in gray). (F) Representative H&E-stained synovial section from an osteoarthritis knee arthroplasty specimen. Red boxes indicate regions containing multinucleated giant cells. (G) Higher-magnification view highlighting a synovial multinucleated giant cell.

Genes positively associated with cartilage degeneration included *PLAUR* (urokinase plasminogen activator receptor), *S100A11*, a RAGE ligand implicated in cartilage catabolism, *CAPG*, linked to macrophage motility and phagocytosis, and *KCNN4*, required for macrophage multinucleation[15,16]. Increased expression of *PRDX1, SLC16A3 (MCT4), HYOU1*, and *ATP5G1* was also observed, consistent with oxidative and metabolic stress in inflamed synovium[17,18]. In contrast, genes negatively associated with cartilage degeneration were enriched for lipid handling and adipocyte-associated metabolic functions, including *DGAT2, FABP4, GPD1, GPAM, SCD, CIDEC, AQP7, LPL, and ADIPOQ*. This pattern is consistent with prior reports showing that adipocyte-associated gene expression in synovium declines with OA progression[11,19].

Visualization of these genes across patient samples demonstrated that synovial transcripts increased in high D-OARSI samples were most strongly expressed in the Lymphomyeloid (C2), Myeloid (C3), and Major trauma (C4) clusters, whereas synovial genes decreased with increased cartilage degeneration were most highly expressed in samples from the Sublining (C1) cluster (Figure 3B). Consistent with these gene-level patterns, pathway analysis revealed that severe cartilage degeneration was associated with enrichment of inflammatory pathways and relative depletion of adipogenesis- and extracellular matrix-associated pathways (Figure 3C). Complete pathway enrichment results are provided in Supplemental Table 10.

We next sought to compare cartilage degeneration-associated bulk RNA-seq genes with synovial cell states. To accomplish this, we integrated our RE-JOIN scRNA-seq data with the previously published AMP-2 synovial scRNA-seq dataset[6] and performed Seurat label transfer. Pseudobulk correlation structure and marker-gene expression across predicted synovial cell states, both globally and within individual lineages and disease diagnoses, are shown in Supplemental Figures 3-4. Per-cell cartilage degeneration gene scores were calculated as the difference between expression of genes positively versus negatively associated with cartilage degeneration, with K-Nearest Neighbor (KNN) smoothing applied to mitigate sparsity-driven noise. Cartilage degeneration scores were highest in lining fibroblasts (median 0.198; IQR 0.160–0.235; n = 1,114) and myeloid cells (median 0.143; IQR 0.114–0.175; n = 9,855), and lowest in sublining fibroblasts (median −0.139; IQR −0.172 to −0.0806; n = 1,496) and endothelial cells (median −0.116; IQR −0.168 to −0.0737; n = 13,250) (Figure 3D). Subtype-level summaries of cartilage degeneration-associated scores are provided in Supplemental Table 4. These findings were consistent with pathway enrichment analyses indicating increased inflammatory and decreased extracellular matrix pathways with cartilage degeneration.

Adipogenesis pathways were preferentially enriched in association with relatively less cartilage degeneration. Because some cell populations, particularly adipocytes and related stromal populations, may be underrepresented in synovial single-cell datasets due to tissue dissociation and capture biases, we next tested whether transcriptional marker genes from a broader set of cell types (Panglao[20]) were associated with cartilage degeneration. Adipocyte-associated GSVA scores were significantly negatively correlated with D-OARSI scores (Spearman ρ = −0.265, P = 0.00197), whereas osteoclast-associated gene set scores were positively correlated with cartilage degeneration severity (ρ = 0.204, P = 0.0183) (Figure 3E). The enrichment of osteoclast-associated gene sets in samples with more severe cartilage degeneration was of interest given our prior observations of multinucleated giant cells in synovium[21,22], and recent reports connecting RANKL inhibition with synovial inflammation and OA progression[23]. In keeping with this pattern, synovial multinucleated giant cells (Figure 3F and 3G) were observed in 21% of samples with severe cartilage degeneration and 7% of samples with less severe degeneration. Together, these results suggest that cartilage degeneration is associated with a loss of adipocyte-linked gene expression and enrichment of inflammatory myeloid features that include osteoclast-associated signals.

### Spatial Transcriptomics Identifies Adipocytes as a Source of Genes Negatively Associated with Cartilage Degeneration

A recent report noted that synovial fibroblasts in normal synovium express adipocyte-associated genes[24]. In our bulk RNA-seq analysis, most genes negatively associated with cartilage degeneration were detectable in scRNA-seq, but many showed low-prevalence and cell-type-restricted expression, limiting our ability to determine whether the “adipocyte-associated” signal reflects fibroblasts or bona fide adipocytes. To resolve the cellular origin of these transcripts, we performed spatial transcriptomics analyses of inflamed OA synovium coupled with paired histology-guided segmentation (Figure 4A,B). After quality filtering, 36,320 cells were retained. Across platforms, the Xenium panel genes were broadly represented in both bulk and single-cell RNA-seq datasets: all 377 genes were present in the scRNA-seq feature space and 77.7% were represented in the bulk RNA-seq expression matrix, with 95.9% showing non-zero expression across bulk samples. At single-cell and spatial resolution, however, gene detection was cell-restricted, with most genes detected in only a minority of cells. The whole-slide cellular composition was dominated by fibroblast populations, including sublining fibroblasts (10,949; 30.1%) and lining fibroblasts (10,582; 29.1%), with additional contributions from dendritic cells (5,620; 15.5%), macrophages (4,447; 12.2%), endothelial cells (2,289; 6.3%), and ACTA2-positive mural cells (1,677; 4.6%) (Figure 4C). Adipocytes were rare across the whole slide (82 of 36,320 cells; 0.23%), consistent with limited capture of adipose-adjacent regions and highlighting a potential blind spot for dissociation-based single-cell datasets.

**Figure 4.**
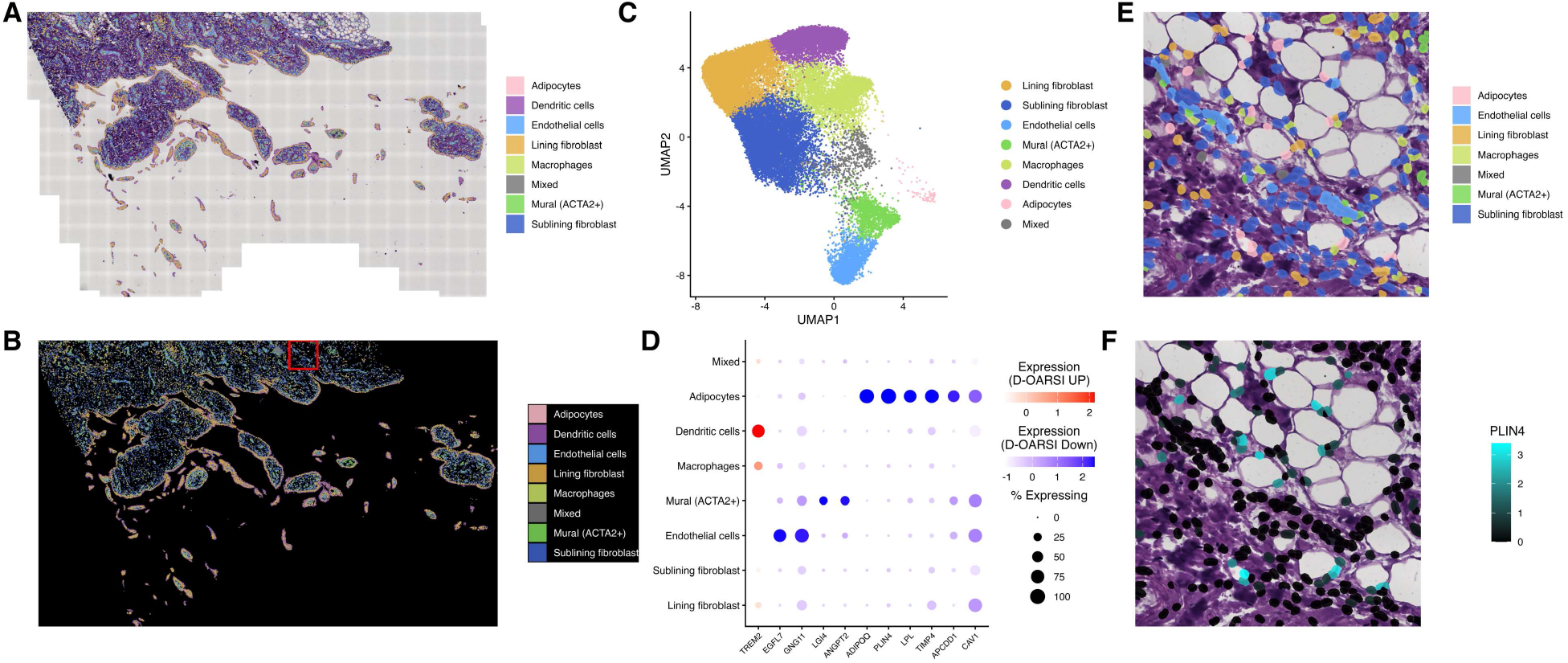
Spatial transcriptomics identifies adipocytes as a source of genes negatively associated with cartilage degeneration. (A,B) Whole-slide OA synovium with single-cell segmentations shown with (A) and without (B) H&E-stained histology overlaid. (C) UMAP of spatial transcriptomic cells colored by annotated synovial cell type, confirming representation of major synovial populations in the spatial dataset. (D) DotPlot of Xenium-captured genes negatively associated with D-OARSI score across synovial cell types. (E) Zoomed view of cell-type segmentations over histology highlighting local tissue organization and adipocyte-rich regions within synovium. (F) Zoomed gene expression projected onto cell segmentations over histology, demonstrating spatial concentration of genes negatively associated with cartilage degeneration within adipocyte-rich areas.

To directly examine adipocyte-driven gene expression, we selected a localized adipocyte-enriched region for higher-resolution inspection (Figure 4E,F). Within this region, adipocytes increased to 17 of 290 cells (5.86%), corresponding to an approximately 26-fold enrichment relative to the whole-slide distribution. This region remained fibroblast-rich, with sublining fibroblasts comprising 162 of 290 cells (55.9%), and also contained endothelial cells (34 of 290; 11.7%), lining fibroblasts (29 of 290; 10.0%), and macrophages (27 of 290; 9.3%), allowing direct comparison of adipocyte-associated transcription with stromal and immune populations in the same tissue context.

We next examined the cellular sources of eleven genes identified by bulk synovial RNA sequencing as associated with cartilage degeneration and captured in the spatial dataset (Figure 4D). Ten genes were negatively associated with D-OARSI and localized predominantly to adipocyte and vascular compartments. Four showed clear adipocyte enrichment by both detection frequency and normalized expression in spatial transcriptomics. *PLIN4* was detected in 100% of adipocytes (mean expression 2.51) but only 4.2% of sublining fibroblasts and 3.4% of lining fibroblasts. *ADIPOQ* was detected in 86.6% of adipocytes (mean 1.29) with minimal detection in sublining fibroblasts (1.08%). *LPL* similarly localized to adipocytes (68.3%, mean 0.762) with markedly lower detection across fibroblast compartments (≤1.1%), while *APCDD1* showed adipocyte-biased detection (57.3%, mean 0.433) with secondary detection in mural (25.5%) and endothelial (21.1%) cells. Several negatively associated genes were enriched in vascular-associated populations. *EGFL7* was most enriched in endothelial cells (69.2%, mean 0.485), whereas *ANGPT2* and *LGI4* showed highest detection in ACTA2-positive mural cells (*ANGPT2*: 31.1%, mean 0.199; *LGI4*: 25.3%, mean 0.145) with lower detection in endothelial cells. Two additional negatively associated genes exhibited broader distribution across stromal and vascular compartments. *CAV1* and *GNG11* were detected at high frequency in both lining fibroblasts and endothelial cells (*CAV1*: 80.4% lining, 80.1% endothelial; *GNG11*: 41.5% lining, 83.3% endothelial), while *TIMP4* showed adipocyte enrichment (78.0%, mean 1.09) but also substantial detection in lining fibroblasts (34.0%). Notably, *TREM2* was the only gene positively associated with D-OARSI and localized primarily to myeloid populations, with strongest detection in dendritic cells (67.0%, mean 0.274) and macrophages (26.8%, mean 0.127), with additional lower-frequency detection in lining fibroblasts (13.2%). These observations suggested preferential detection of several negatively associated transcripts in adipocytes compared with other synovial cell populations, prompting formal evaluation of spatial localization.

To test whether these transcripts exhibited non-random spatial localization, we compared the fraction of gene-positive cells within the adipocyte-rich zoom region to the remainder of the tissue using Fisher’s exact testing with false discovery rate correction. Canonical adipocyte-associated transcripts showed strong spatial enrichment, with *ADIPOQ* and *PLIN4* demonstrating approximately 12.5-fold and 7.7-fold enrichment of gene-positive cells within the zoom region, respectively, while *LPL* showed more moderate enrichment of approximately 4.0-fold. Vascular-associated transcripts showed weaker or mixed spatial effects: *EGFL7* and *LGI4* were modestly enriched (approximately two-fold), whereas broadly expressed genes such as *CAV1* and *GNG11* were slightly depleted, and *ANGPT2, TIMP4*, and *APCDD1* showed no significant localization bias after correction. Together, these findings indicate that bulk synovial decreased adipocyte-associated transcripts in bulk synovium likely reflect loss of fat-rich architecture and increased inflammatory/fibrotic remodeling, with additional contributions from vascular cell populations, while spatial profiling resolves a distinct myeloid signal for the single positively associated gene *TREM2*.

## Discussion

Recent transcriptomics analyses have highlighted heterogeneity in human OA synovium, which segregates into two-to-four molecular phenotypes [7–9]. Imaging studies have demonstrated that synovitis correlates with cartilage degeneration and predicts progressive cartilage loss in OA [12–14]. However, the molecular features linking synovial heterogeneity to cartilage degeneration remain incompletely defined. Here, we identify four synovial transcriptional clusters in human knee OA that differ in histologic features and cartilage degeneration severity. Clusters associated with more severe degeneration were enriched for lining fibroblast- and inflammatory myeloid-associated gene expression, whereas the least severe cluster (sublining, C1) showed enrichment of sublining fibroblast, endothelial, and mural cell markers.

Comparison of bulk and single-cell RNA-seq enabled cell-type interpretation and biological context of these clusters. We defined sublining (C1), lymphomyeloid (C2), and myeloid (C3) clusters, while C4 was enriched in patients reporting prior major trauma to the operated knee, suggesting injury-associated synovial remodeling as a contributor to molecular heterogeneity. Given the small sample size, particularly in C4, these findings require replication in larger cohorts.

Genes associated with cartilage degeneration reflected inflammatory, macrophage, and osteoclast-associated gene expression. The presence of multinucleated giant cells, together with increased expression of *KCNN4*[15], suggests multinucleated inflammatory myeloid states in OA synovium. Prior work implicates OSCAR signaling in cartilage destruction [25] and TREM2-DAP12 signaling in osteoclast differentiation [26,27], raising the possibility that TREM2-expressing synovial myeloid populations may adopt osteoclast-like or multinucleated inflammatory phenotypes, although this will require direct validation.

In contrast, genes negatively associated with cartilage degeneration were enriched for adipocyte-associated and lipid-handling transcripts. Spatial transcriptomic profiling demonstrated that canonical adipocyte-associated transcripts localized predominantly to adipocytes, with additional contributions from vascular and stromal compartments. These signals may mark synovial regions with preserved adipocyte-rich and stromal-vascular architecture and reduced inflammatory remodeling, consistent with prior reports demonstrating loss of adipocytes and adipocyte-associated gene expression in OA synovium[19,24,28] as well as a decrease in adipose tissue area, adipocyte size, and expression of *PLIN* in rabbits after anterior cruciate ligament transection and partial medial menisectomy[29].

Several limitations should be considered. The cross-sectional design precludes causal inference, and bulk RNA-seq may confound changes in cell state and composition. Synovial sampling was based on grossly apparent disease rather than standardized anatomical mapping, and biopsy location was not uniformly recorded. Spatial analyses were performed in a single specimen and should be considered hypothesis-generating. Finally, these samples were derived from patients undergoing total knee arthroplasty, limiting generalizability to earlier OA.

Despite these limitations, our results support a model in which OA comprises biologically distinct synovial clusters linked to cartilage degeneration severity. By linking synovial gene expression to cartilage degeneration and organizing these signals into molecular subtypes, this work provides a framework for stratifying OA and prioritizing synovial pathways for mechanistic and therapeutic investigation.

## Ethics approval and consent to participate

All procedures were performed in accordance with the Declaration of Helsinki. Human biospecimens used in this study were obtained under protocols approved by the Institutional Review Boards of The Rockefeller University (DOR-0822) and the Hospital for Special Surgery (2014-233). Written informed consent was obtained from all participants prior to inclusion in the study. Material transfer agreements were in place between institutions for the sharing and analysis of biospecimens.

## Data availability

The data that support the findings of this study are available from the Accelerating Medicines Partnership (AMP) RA/SLE program via the ARK Portal (https://arkportal.synapse.org/). Additional data generated or analyzed during this study are available from the corresponding author upon reasonable request, subject to institutional and data use agreements.

## Acknowledgements

The authors acknowledge the Tow Foundation for support of the David Z. Rosensweig Genomics Research Center. Technical support was provided by the HSS Molecular Histopathology Core Laboratory.

## The RE-JOIN consortium consists of

Armen Akopian, Kyle Allen, Alejandro Almarza, Benjamin Arenkiel, Yangjin Bae, Bruna Balbino de Paula, Anita Bandrowski, Mario Danilo Boada, Jacqueline Boccanfuso, Jyl Boline, Dawen Cai, Dellina Lane, Robert Caudle, Racel Cela, Yong Chen, Rui Chen, Brian Constantinescu, Yenisel Cruz-Almeida, M. Franklin Dolwick, Chris Donnelly, Zelong Dou, Joshua Emrick, Malin Ernberg, Danielle Freburg-Hoffmeister, Spencer Fullam, Janak Gaire, Akash Gandhi, Benjamin Goolsby, Stacey Greene, Nele Haelterman, Michael Iadarola, Shingo Ishihara, Azeez Ishola, Sudhish Jayachandran, Zixue Jin, Frank Ko, Priya Kulkarni, Zhao Lai, Brendan Lee, Yona Levites, Carolina Leynes, Jun Li, Martin Lotz, Lindsey Macpherson, Tristan Maerz, Camilla Majano, Anne-Marie Malfait, Maryann Martone, Bella Mehta, Richard Miller, Rachel Miller, Michael Newton, Alia Obeidat, Merissa Olmer, Dana E. Orange, Miguel Otero, Kevin Otto, Folly Patterson, Marlena Pela, Sienna Perry, Theodore Price, Hernan Prieto, Russell Ray, Dongjun Ren, Margarete Ribeiro Dasilva, Alexus Roberts, Elizabeth Ronan, Oscar Ruiz, Shad Smith, Mairobys Socorro, Kaitlin Southern, Joshua Stover, Michael Strinden, Hannah Swahn, Evelyne Tantry, Sue Tappan, Luis Tovias Sanchez, Airam Vivanco-Estela, Joost Wagenaar, Lai Wang, Kim Worley, Joshua Wythe, Jiansen Yan, and Julia Younis.

## The Accelerating Medicines Partnership: RA/SLE Network (AMP RA/SLE) consists of

Fan Zhang, Anna Helena Jonsson, Aparna Nathan, Nghia Millard, Michelle Curtis, Qian Xiao, Maria Gutierrez-Arcelus, William Apruzzese, Gerald F. M. Watts, Dana Weisenfeld, Saba Nayar, Javier Rangel-Moreno, Nida Meednu, Kathryne E. Marks, Ian Mantel, Joyce B. Kang, Laurie Rumker, Joseph Mears, Kamil Slowikowski, Kathryn Weinand, Dana E. Orange, Laura Geraldino-Pardilla, Kevin D. Deane, Darren Tabechian, Arnoldas Ceponis, Gary S. Firestein, Mark Maybury, Ilfita Sahbudin, Ami Ben-Artzi, Arthur M. Mandelin II, Alessandra Nerviani, Myles J. Lewis, Felice Rivellese, Costantino Pitzalis, Laura B. Hughes, Diane Horowitz, Edward DiCarlo, Ellen M. Gravallese, Brendan F. Boyce, Larry W. Moreland, Susan M. Goodman, Harris Perlman, V. Michael Holers, Katherine P. Liao, Andrew Filer, Vivian P. Bykerk, Kevin Wei, Deepak A. Rao, Laura T. Donlin, Jennifer H. Anolik, Michael B. Brenner, Soumya Raychaudhuri

