## Supplemental Materials for "Synovial transcriptional clusters link cartilage degeneration to cell-type-specific gene expression in knee osteoarthritis"

### Supplemental Legends

### Figures

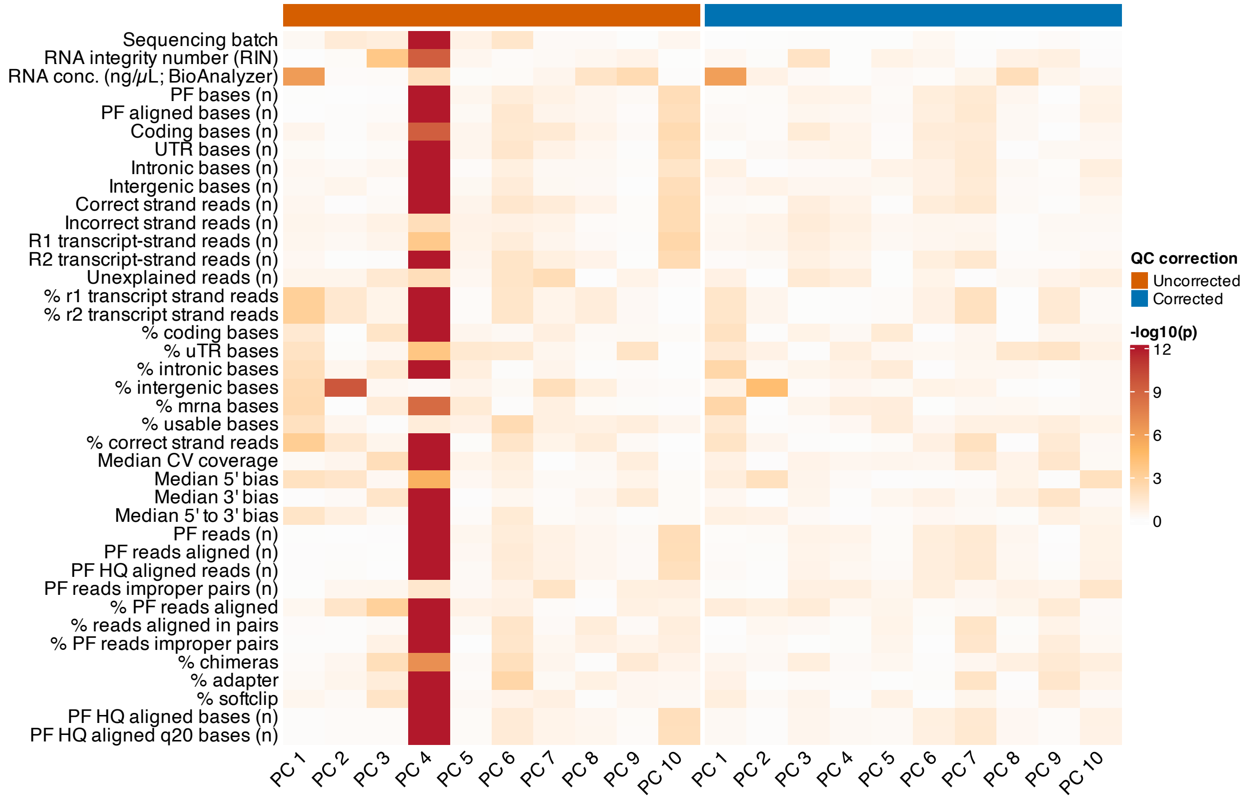


**Supplemental Figure 1. Adjustment of RNA expression matrix by batch and intergenic reads reduces associations between principal components and technical QC metrics.** Heatmap showing associations between principal components (PC1–PC10) derived from bulk synovial RNA-seq data and sequencing quality control (QC) metrics. Values represent −log10(p-values) from correlations between each QC metric and individual PCs. Columns are split by expression matrix before correction (Uncorrected, left) and after adjustment for sequencing batch and percent intergenic reads (Corrected, right). QC metrics include sequencing batch, total reads, percent aligned reads, transcriptomic composition metrics (percent coding, intronic, intergenic, and mRNA bases), and median 3′ bias. Reduced signal following correction indicates decreased contribution of technical variation to major axes of transcriptomic variance.

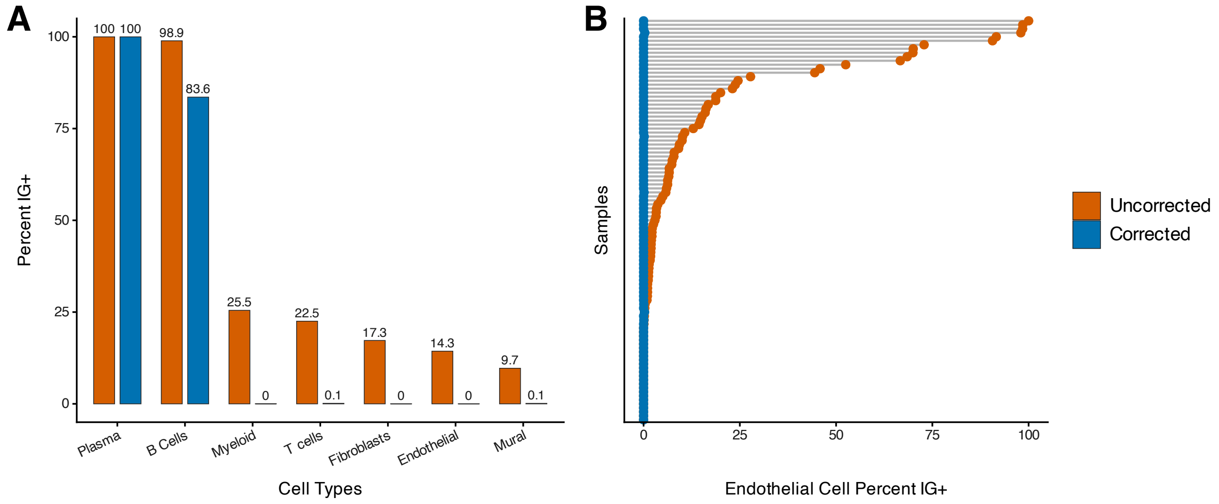


**Supplemental Figure 2. Representative example showing reduction of immunoglobulin gene spillover into non-B lineage cell types following iterative two-stage DecontX correction.** (A) Percent IG+ cells by cell type comparing expression before (Uncorrected) and after correction (Corrected). Cells were classified as IG+ based on immunoglobulin gene expression. (B) For endothelial cells, percent IG+ cells per sample are shown before and after correction, with samples ordered by the uncorrected value.

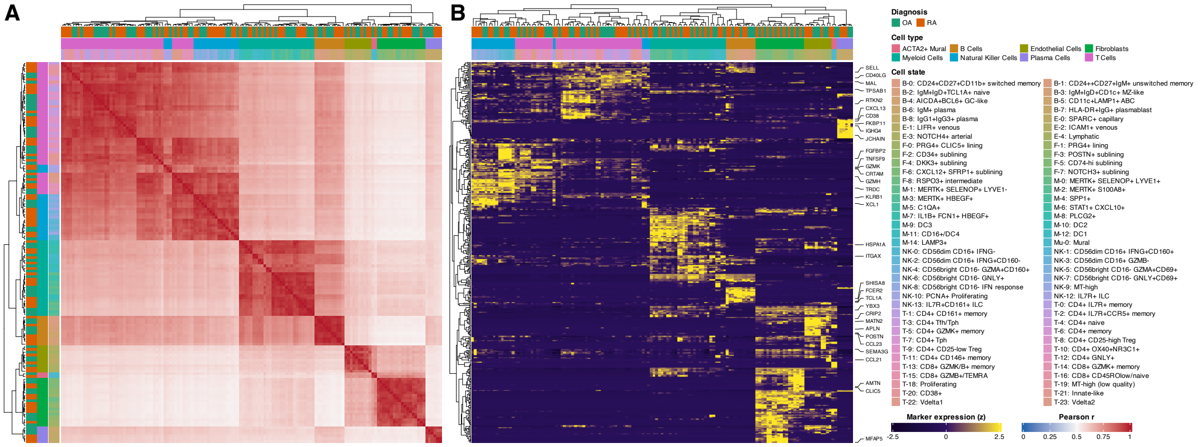


**Supplemental Figure 3. Global pseudobulk similarity and marker-gene expression across synovial cell states.** (A) Pearson correlation heatmap between pseudobulk expression computed using the top 3,000 most variable genes across all cell types. Each row and column correspond to a bucket defined by diagnosis (OA vs RA) and predicted subtype/cell state (minimum 10 cells per bucket). Correlations are displayed on a 0–1 scale. (B) Heatmap of published AMP-2 marker-gene pseudobulk expression across the same buckets, using the union of marker genes across cell types. Values show row-wise z-scored pseudobulk expression, with selected marker genes labeled. Top annotations indicate diagnosis, cell type, and subtype/cell state.

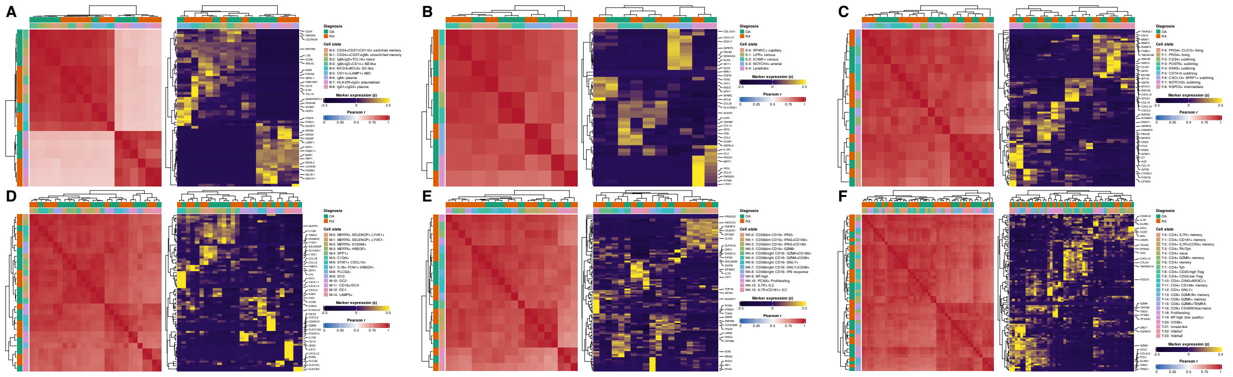

**Supplemental Figure 4. Cell-type–specific pseudobulk similarity and marker-gene expression across synovial cell states.** For each cell type, Pearson correlation heatmaps between pseudobulk expression computed using the top 3,000 most variable genes across buckets defined by diagnosis (OA vs RA) and subtype/cell state (minimum 10 cells per bucket), displayed on a 0–1 scale (left). Marker-gene heatmaps show pseudobulk expression across the same buckets, displayed as row-wise z-scored expression with selected marker genes labeled (right). (A) Endothelial cells. (B) Fibroblasts. (C) Myeloid cells. (D) B lineage cells. (E) T cells. (F) Natural killer cells.

### Tables

**Supplemental Table 1. Bulk RNA-seq technical quality control metrics for osteoarthritis synovial samples.** The table reports per-sample sequencing and alignment quality metrics derived from Picard RNA-seq and alignment analyses, together with RNA integrity number (RIN) and BioAnalyzer RNA concentration measurements. Metrics include read and base counts, genomic feature distribution (coding, UTR, intronic, and intergenic bases), strand specificity measures, coverage and transcript bias statistics, and alignment performance metrics. Original Picard metric column names are retained.

**Supplemental Table 2. Differential gene expression associated with cartilage degeneration severity (D-OARSI High vs Low).** Differential expression was tested using bulk RNA-seq counts with voom+limma, comparing samples with high cartilage degeneration (D-OARSI score ≥20) versus low degeneration (D-OARSI score <20). Models included sequencing batch and percent intergenic bases as covariates. The table reports log2 fold change (High vs Low), t-statistic, average expression, P value, and FDR-adjusted P value (BH). Positive log2 fold change indicates higher expression in the High group.

**Supplemental Table 3. Sample-level cell tracking through single-cell RNA-seq preprocessing in osteoarthritis synovium.** For each osteoarthritis synovial sample in the RE-JOIN cohort, the table reports the number of cells retained at successive stages of single-cell RNA-seq preprocessing, including initial cell detection, quality control filtering, and doublet removal. Columns correspond to cell counts after each processing step, allowing tracking of cell retention across the analysis workflow. Hashed and multiplexed samples are indicated where applicable, and samples excluded because of excessive cell loss are noted. Batch processing dates are provided for reference.

**Supplemental Table 4. Single-cell marker genes by cell type in osteoarthritis synovium.** Marker genes were identified for each cell type in osteoarthritis synovium (RE-JOIN cohort) by comparing each cell type with all others using a Wilcoxon rank-sum test. Genes were retained based on expression enrichment and prevalence within the target cell type, with common housekeeping, mitochondrial, ribosomal, HLA, and most immunoglobulin genes excluded to avoid dominance by broadly expressed transcripts. For Figure 2 visualizations, the top 30 markers per cell type were selected after filtering and restricted to genes present in the bulk RNA-seq PCA gene set (top 5,000 genes by MAD). The table reports log2 fold-change, AUC, P value, FDR-adjusted P value, and the fraction of cells expressing each gene inside and outside the listed cell type.

**Supplemental Table 5. Clinical comparability of bulk and single-cell osteoarthritis cohorts.** Clinical and demographic characteristics of individuals with knee osteoarthritis included in bulk transcriptomic and single-cell transcriptomic cohorts are shown. Continuous variables are reported as mean (SD) when normally distributed, as assessed by the Shapiro–Wilk test, and as median (p25, p75) otherwise; categorical variables are reported as n (%). P values were calculated using Wilcoxon rank-sum tests for continuous variables and Fisher's exact tests for categorical variables. Statistically significant values ($P < 0.05$) are shown in bold.

**Supplemental Table 6. Differentially expressed genes for each transcriptional cluster compared with all other clusters.** For each cluster (C1–C4), gene expression was compared with all other clusters combined. The table is filtered to include only genes with positive log2 fold-change (log2FC), indicating higher expression in the listed cluster. P values and FDR values indicate statistical significance.

**Supplemental Table 7. Differentially expressed genes for each transcriptional cluster compared with cluster C1.** For clusters C2–C4, gene expression was compared with cluster C1 using limma-voom. The table is filtered to include only genes with positive log2 fold-change (log2FC), indicating higher expression in the listed cluster. P values and FDR values indicate statistical significance.

**Supplemental Table 8. Pathway enrichment for each transcriptional cluster compared with all other clusters.** For each cluster (C1–C4), pathway enrichment was tested using genes ranked by their association with that cluster compared with all others. Positive NES values indicate pathways enriched in the listed cluster. FDR values indicate statistical significance, and leading-edge genes show the genes contributing most to enrichment.

**Supplemental Table 9. Pathway enrichment for each transcriptional cluster compared with cluster C1.** For clusters C2–C4, pathway enrichment was tested using genes ranked by their association with that cluster compared with cluster C1. Positive NES values indicate pathways enriched in the listed cluster. FDR values indicate statistical significance, and leading-edge genes show the genes contributing most to enrichment.

**Supplemental Table 10. Pathway enrichment associated with cartilage degeneration severity (D-OARSI High vs Low).** Pathway enrichment was tested using FGSEA on MSigDB Hallmark and Reactome gene sets, ranking genes by the differential expression effect (log2 fold change) from Supplemental Table 2 (High vs Low). The table reports normalized enrichment score (NES), P value, and FDR-adjusted P value (BH). Positive NES indicates pathways enriched in the High group, and negative NES indicates pathways enriched in the Low group.

**Supplemental Table 11. Cartilage degeneration–associated gene expression by synovial cell subtype in osteoarthritis.** For each synovial cell state, the table reports the number of cells and summary statistics (mean and median) for single-cell scores derived from genes associated with higher cartilage degeneration, lower cartilage degeneration, and their difference in osteoarthritis synovium (RE-JOIN cohort). Only subtypes with at least 50 cells are included.

### Methods

#### Patient Cohort

A total of n = 154 patients with knee OA undergoing total knee arthroplasty (TKA) were enrolled from a convenience sample at a high-volume tertiary care hospital approximately two weeks prior to surgery, with Institutional Review Board approval (IRB #2018-0895) and patient consent. Synovial tissues from n = 135 patients were used for bulk RNA-seq. Additional synovial tissues were processed for single-cell RNA-seq (scRNA-seq, n = 18) and spatial transcriptomics (n = 1). Patients over 45 years of age undergoing TKA who met the 1986 American College of Rheumatology (ACR) Clinical and Radiographic Criteria[1], ACR Clinical and Laboratory Criteria, and Kellgren-Lawrence (KL) radiographic criteria[2] (grades 2–4) for knee OA were included. Patients with a prior TKA, fracture of the operative knee, inflammatory or autoimmune rheumatic disease (including rheumatoid arthritis, spondyloarthropathy, ankylosing spondylitis, or psoriatic arthritis), or indications for TKA other than OA were excluded. Clinical characteristics including age, sex, race, body mass index (BMI), duration of symptoms, duration since diagnosis, pain duration, erythrocyte sedimentation rate (ESR), and high-sensitivity C-reactive protein (hsCRP) were collected. KL grades were assessed by a musculoskeletal radiologist using preoperative knee radiographs. The study was approved by the HSS Institutional Review Board (IRB #2018-0895) and the Rockefeller University Review Board (#DOR0822).

#### Tissue Retrieval, Processing, and Histologic Scoring

Joint tissues retrieved at the time of TKA were immediately transported to the HSS Department of Pathology and Laboratory Medicine for processing and selection. For each of the first n = 135 patients enrolled in the study, an expert musculoskeletal pathologist (E.D.) grossly inspected the synovium and selected the most obviously diseased region for biopsy, defined by features such as hyperplastic papillary architecture, dull or opaque appearance, and obscuration of the underlying vasculature. Synovial tissue fragments were then preserved as formalin-fixed, paraffin-embedded (FFPE) tissue for histologic analyses, with an adjacent sample preserved in RNAlater for RNA isolation and bulk RNA-seq. Cryopreserved (CryoStor® CS10) synovial tissues retrieved from a subsequent subset of patients (n = 18) were used for viable cell isolation and scRNA-seq. FFPE-preserved synovium (n = 1) was used for Xenium spatial transcriptomics. Osteochondral slabs selected from the most severely eroded and relatively preserved condyles according to gross inspection of the first n = 135 patients were formalin-fixed, decalcified, processed through a standard cycle, and paraffin-embedded.

Cartilage histopathology was scored by an expert musculoskeletal pathologist (E.D.) using the Osteoarthritis Research Society International (OARSI) grading system[3]. Scores were assessed separately on 5-μm-thick sections from the most severely eroded femoral condyle and from a relatively preserved condyle according to gross inspection. For downstream analyses, the OARSI score from the most severely eroded condyle was used as the primary quantitative measure of cartilage degeneration (D-OARSI). A threshold of D-OARSI ≥ 20 was selected to distinguish samples with more advanced cartilage degeneration within this arthroplasty cohort, in which overall histopathologic severity was high.

Synovial histology was evaluated on 5-μm-thick H&E-stained sections and scored for ten features: lymphocytic inflammation, lining hyperplasia, binucleate plasma cells, Russell bodies, synovial giant cells, fibrin, fibrosis, synovial mucoid degeneration, detritus, and whether plasma cells constituted more than 10% of lymphocytes, as previously described[4]. Additional synovial inflammation metrics, including cell density and cellular aggregates, were quantified using a previously developed computer vision-based algorithm[5].

### Synovial Tissue RNA Isolation and Bulk RNA Sequencing

For bulk RNA-seq, RNA was isolated from RNAlater-preserved synovial tissue fragments using TRIzol followed by DNase I digestion and column cleanup (Qiagen). RNA quantity and integrity were assessed using a Bioanalyzer. cDNA libraries were prepared using the KAPA Stranded RNA-Seq Kit with RiboErase and sequenced on a NovaSeq platform (150-bp paired-end reads), targeting approximately 100 million reads per sample.

### Bioinformatic Analyses of Bulk RNA Sequencing

Sequencing quality control metrics were assessed using Picard[6] (v2.27.5) tools and are summarized in Supplemental Table 1. Transcript abundances were quantified using kallisto[7] (v0.46.2) and summarized to gene-level counts and abundance estimates using tximport[8] (v1.30.0) with transcript-to-gene mappings based on Ensembl annotations. Genes were restricted to protein-coding and immunoglobulin loci, and entries were collapsed by Ensembl gene ID across samples. Normalized log counts per million (logCPM) values were generated using edgeR[9] (v4.4.1) following library size normalization.

Associations between Picard-derived sequencing quality metrics and principal components of gene expression were evaluated to identify technical sources of variation. Because percent intergenic bases correlated with a major axis of transcriptional variation (Supplemental Figure 1), expression matrices were adjusted for percent intergenic bases and sequencing batch using limma’s removeBatchEffect() prior to consensus clustering.

Differential expression analyses comparing samples with high versus low cartilage degeneration severity defined by D-OARSI thresholds were performed using limma-voom linear modeling with sequencing batch and percent intergenic bases included as covariates in the design matrix (Supplemental Table 2). Gene set enrichment analysis was performed using fgsea[10] (v1.30.0) on ranked limma-voom[11,12] (limma v3.60.6) differential expression statistics using MSigDB Hallmark[13] and Reactome[14] pathway gene sets. Cell-type–associated enrichment analyses were conducted using gene sets derived from the Panglao Cell Atlas[15]. Gene set variation analysis[16] (GSVA (v1.52.3)) was used to compute sample-level enrichment scores for selected gene sets, which were correlated with cartilage degeneration severity. Downstream analyses were performed in R (v4.4.1; Bioconductor v3.19).

### Single-Cell RNA Sequencing

### Synovial Tissue Dissociation and Sequencing

Cryopreserved human OA synovial tissues from the RE-JOIN osteoarthritis cohort (n = 18 patients) were dissociated into single-cell suspensions using a combined enzymatic and mechanical protocol adapted from the Accelerating Medicines Partnership (AMP) Rheumatoid Arthritis and Systemic Lupus Erythematosus (AMP RA/SLE) Network[17]. Briefly, tissue fragments were rapidly thawed and digested in RPMI containing Liberase™ TL (100 µg/mL; Roche, Cat. No. 05401020001) and DNase I (100 µg/mL; Roche, Cat. No. 10104159001) for 30 minutes at 37°C in a MACS rotator. An additional treatment with Trypsin-EDTA (0.25%) with phenol red (Gibco, Cat. No. 25200056) was performed at room temperature for 5 minutes. Following digestion, samples were mechanically dissociated and filtered, and viable cells were isolated by fluorescence-activated cell sorting (FACS) prior to scRNA-seq.

### Single-cell RNA Sequencing Data Processing and Quality Control

10x Genomics single-cell RNA sequencing reads were aligned to the GRCh38 human genome (2020 reference) using CellRanger[18] (v8.0.0), generating gene-cell count matrices across sequencing runs. Across 28 sequencing libraries (19 runs), 170,138 cells were initially recovered prior to quality control filtering. Samples were demultiplexed using Seurat[19] (v5.3.0) HTODemux()[20] with default parameters. For samples sequenced jointly without hashing, demultiplexing was performed using genotype-based approaches implemented in Vireo[21] (v0.5.8) and Souporcell[22] (v2.4), with sample identity supported by expression of sex-linked genes. Doublets were identified independently within each dataset using DoubletFinder[23] (v2.0.6) and removed prior to downstream analysis.

Each sample was processed individually by estimating mitochondrial transcript content and applying quality control filters to retain cells with 500-5,000 detected genes and mitochondrial transcript fractions ≤20%, thereby excluding low-quality droplets and likely multiplets (Supplemental Table 3). Unique molecular identifier (UMI) counts were evaluated during quality control but were not subjected to a fixed threshold beyond these feature-based filters. Samples in which more than 40% of initially recovered cells were removed during demultiplexing and quality control filtering were excluded from downstream analyses, consistent with quality control criteria established by AMP RA/SLE consortium analyses. After filtering, 124,465 cells RE-JOIN osteoarthritis synovial samples were retained for downstream analysis.

**Ambient RNA Correction of Single-cell Transcriptomes**

Ambient RNA contamination was corrected using DecontX[24] (v1.4.1) in a two-stage framework following the recommended iterative DecontX workflow. DecontX estimates per-cell contaminant fractions and contamination-adjusted expression profiles using a Bayesian mixture model. First, DecontX was applied in an unsupervised manner using empty-droplet–derived background signal to model experiment-wide ambient RNA contamination. Residual expression of cell-type-restricted transcripts across multiple cell populations after the initial pass motivated a second application incorporating broad cell-type annotations (lymphoid, myeloid, fibroblast, endothelial, and mural compartments) to guide estimation of cross-compartment contamination estimation. B cells and plasma cells were merged to model immunoglobulin-producing populations during this step (Supplemental Figure 2). Corrected counts from the second pass were stored as a parallel assay and log-normalized for downstream gene expression analyses.

**AMP-2 and RE-JOIN Dataset Integration, Clustering, and Reference-based Label Transfer**

For integration with external AMP: RA/SLE Network (AMP) osteoarthritis and rheumatoid arthritis synovial datasets[25], the top 3,000 variable features were selected, followed by principal component analysis (PCA). Batch effects across samples were corrected using Harmony[26] (v1.2.3), after which neighbor graphs and UMAP embeddings were computed for downstream clustering and visualization. Clustering was performed using Seurat[19], with resolution selected based on cluster stability analysis using Clustree[27] to minimize lineage mixing while preserving biologically coherent populations. Cell state labels were assigned by reference-based mapping to AMP-2 using Seurat FindTransferAnchors() and TransferData(). Integration and annotation consistency across datasets were evaluated using pseudobulk correlation and marker-gene analyses (Supplemental Figures 3–4).

**Marker Gene Identification and Construction of Cell-Type Gene Sets**

Cell-type marker genes were identified from synovial scRNA-seq data following ambient RNA correction. Cells were grouped according to assigned cell-type identities based on canonical lineage markers[25], and differential expression analysis was performed by comparing each cell type with all others. Marker discovery was conducted using a Wilcoxon rank-sum framework implemented in the presto[28] package. For each gene and cell type, effect size and discrimination metrics were calculated, including $\log_2$ fold-change, area under the receiver operating characteristic curve (AUC), and the fraction of cells expressing the gene within the target population relative to other cell types.

Marker genes were filtered to retain robust cell-type-enriched signals, requiring $\log_2$ fold-change ≥0.5, expression in ≥40% of cells within the target cell type, and detection in at least 10% more cells than in the remaining dataset. Ribosomal, mitochondrial, and selected housekeeping genes were excluded to reduce broadly expressed or technical signals, and *HLA* genes were excluded where specified. Immunoglobulin genes were generally excluded from plasma-cell markers to prevent dominance by antibody transcripts while preserving representative lineage-defining genes. The resulting marker gene sets are provided in Supplemental Table 4.

To relate bulk transcriptional variation to cell-type-associated expression, PCA was performed on bulk synovial RNA-seq data, and gene loadings for PC1 and PC2 were extracted. For each major synovial cell type, top-ranked marker genes were intersected with genes present in the bulk PCA gene set, and distributions of PC loadings were visualized to assess alignment of cell-type-associated genes with dominant bulk transcriptional axes.

**Xenium Spatial Transcriptomics Profiling**

Spatial transcriptomics profiling of OA synovium was performed using the Xenium In Situ Platform (10x Genomics) on a single FFPE synovial tissue specimen obtained at the time of TKA. Briefly, synovium was immediately fixed in freshly-prepared 10% formalin for 24 hours, washed in 1X PBS, and paraffin-embedded following standard procedures. The resulting FFPE blocks were stored at 4°C until sectioned for downstream analyses.  During FFPE handling, all surfaces, microtome blades and sectioning tools were cleaned with RNaseZap™ RNase Decontamination Solution (Invitrogen Catalog # AM9780), and nuclease-free water (ThermoFisher Catalog # 4387936) was used to float sections in a water bath maintained at 50°C.

Before analyses, we conducted a series of quality control (QC) steps. First, we calculated the DV200 value from RNA isolated from FFPE sections (15-µm-thick total) and analyzed using a Bioanalyzer (Weill Cornell Genomics Resources Core Facility). All samples analyzed had DV200 scores > 30%, meeting the threshold recommended by 10x Genomics for optimal Xenium analyses. Next, we obtained and H&E-stained one 5-µm section, which was evaluated by a pathologist (D.R.) to confirm appropriate orientation and tissue quality. One adjacent 5-µm paraffin section was mounted in the capture area of a Xenium slide (equilibrated to room temperature for at least 30 min) without covering the fiducials and used for downstream analysis according to the manufacturer's instructions (Xenium In Situ Gene Expression # CG000582 protocol for Xenium V1 platform).

A pre-designed panel with n = 377 genes (V1 Xenium Human Multi-Tissue and Cancer Panel; Catalog # 1000626) was selected because it includes genes that are expressed in multiple cells and tissue types. The Xenium Analyzer (v2.2.0.1) was used for imaging and transcript detection. Exported transcript and segmentation outputs were processed and analyzed in R using Seurat v5 and associated spatial transcriptomics workflows.

Cell segmentation was based on nuclear staining with morphological expansion to approximate cellular boundaries, generating per-cell transcript counts and spatial coordinates. Cells with low transcript counts or segmentation artifacts were excluded during QC. Resulting gene-cell count matrices and spatial coordinates were analyzed in R for downstream analyses.

Cell-type annotation was performed by comparing per-cell gene expression profiles with marker gene sets derived from synovial scRNA-seq, enabling classification of major synovial cell populations. Region-of-interest analyses were subsequently performed using histologic morphology and spatial gene expression patterns to evaluate localization of transcripts within regions with intact synovial architecture.

**Statistical Analysis**

Statistical analyses were performed in R (v4.4.1). For comparisons of continuous variables across more than two groups, Welch's ANOVA or Kruskal-Wallis tests were used as appropriate based on distributional assumptions. Post hoc comparisons were performed using Dunnett's tests when comparing clusters to a reference group. For pairwise comparisons, Wilcoxon rank-sum tests were used. One-sample Wilcoxon tests were used to assess whether cluster-level gene expression enrichment scores differed from zero. Associations between categorical variables were evaluated using Fisher's exact tests. Correlations between continuous variables were assessed using Spearman's rank correlation coefficients. Where applicable, P values were adjusted for multiple testing using the Benjamini-Hochberg false discovery rate method. Statistical significance was defined as a two-sided $P$ value $< 0.05$.
